# Red blood cell passage through deformable interendothelial slits in the spleen: Insights into splenic filtration and hemodynamics

**DOI:** 10.1101/2024.02.22.581664

**Authors:** Guansheng Li, He Li, Papa Alioune Ndou, Mélanie Franco, Yuhao Qiang, Xuejin Li, Pierre A. Buffet, Ming Dao, George Em Karniadakis

## Abstract

The spleen constantly clears altered red blood cells (RBCs) from the circulation, tuning the balance between RBC formation (erythropoiesis) and removal. The retention and elimination of RBCs occur predominantly in the open circulation of the spleen, where RBCs must cross submicron-wide inter-endothelial slits (IES). Several experimental and computational studies have illustrated the role of IES in filtrating the biomechanically and morphologically altered RBCs based on a rigid wall assumption. However, these studies also reported that when the size of IES is close to the lower end of clinically observed sizes (less than 0.5 *μ*m), an unphysiologically large pressure difference across the IES is required to drive the passage of normal RBCs, sparking debates on the feasibility of the rigid wall assumption. In this work, we perform a computational investigation based on dissipative particle dynamics (DPD) to explore the impact of the deformability of IES on the filtration function of the spleen. We simulate two deformable IES models, namely the passive model and the active model. In the passive model, we implement the worm-like string model to depict the IES’s deformation as it interacts with blood plasma and allows RBC to traverse. In contrast, the active model involved regulating the IES deformation based on the local pressure surrounding the slit. To demonstrate the validity of the deformable model, we simulate the filtration of RBCs with varied size and stiffness by IES under three scenarios: 1) a single RBC traversing a single slit; 2) a suspension of RBCs traversing an array of slits, mimicking *in vitro* spleen-on-a-chip experiments; 3) RBC suspension passing through the 3D spleen filtration unit known as ‘the splenon’. Our simulation results of RBC passing through a single slit show that the deformable IES model offers more accurate predictions of the critical cell surface area to volume ratio that dictate the removal of aged RBCs from circulation compared to prior rigid-wall models. Our biophysical models of the spleen-on-a-chip indicates a hierarchy of filtration function stringency: rigid model > passive model > active model, providing a possible explanation of why the spleen-on-a-chip could overestimate the filtration function of IES. We also illustrate that the biophysical model of ‘the splenon’ enables us to replicate the *ex vivo* experiments involving spleen filtration of malaria-infected RBCs. Taken together, our simulation findings indicate that the deformable IES model could serve as a mesoscopic representation of spleen filtration function closer to physiological reality, addressing questions beyond the scope of current experimental and computational models and enhancing our understanding of the fundamental flow dynamics and mechanical clearance processes within in the human spleen.

## 1. Introduction

The spleen, positioned in the abdomen near the greater curvature of the stomach [1], stands as the largest secondary immune organ in humans. It orchestrates immune responses to halt the proliferation of pathogenic microorganisms within the bloodstream and acts as a vital blood filter to eliminate aged, damaged, and diseased blood cells from circulation [2, 3]. This dual functionality is achieved through its distinct compartments, the white and red pulp [4, 2]. Comprising T- and B-cells encircling arterial vessels, the white pulp executes immune functions, while the red pulp, characterized by venous sinuses and a reticular meshwork of splenic cords housing macrophages, serves as a blood filter [5, 6, 7, 3]. Within the red pulp, red blood cells (RBCs) undergo meticulous scrutiny by macrophages, navigate through narrow interendothelial slits (IES), and return to venous sinuses, a process crucial for maintaining the quality of circulating RBCs by mechanically retaining less deformable ones and facilitating their phagocytosis by macrophages [8, 9, 10]. Approximately 90% of the overall blood flow within the spleen circumvents the reticular meshwork in the red pulp [11], instead proceeding directly towards the neighboring venous sinuses, a phenomenon referred to as the “fast pathway.” Conversely, the remaining approximately 10% of the blood takes the “slow pathway,” entering the reticular meshwork within the red pulp before progressing further [12, 3]. This complex interplay of biochemical and biomechanical processes is pivotal for the spleen’s continuous quality control over circulating RBCs, underscoring its indispensable role in the immune and circulatory systems of the body.

The sinusoidal network in the spleen comprises endothelial cells aligned in parallel and interconnected by stress fibers to annular fibers composed of extracellular matrix constituents. Characterized by a narrower and shorter configuration than capillaries, IES induce a dumbbell-shaped deformation in RBCs during their transit [13]. This unique three-dimensional process has only been directly observed in rodent models [12], facilitated by the visualization of their spleens. Comprehensive investigations into the mechanisms governing the clearance of normal and diseased RBCs by the human spleen face two primary challenges [14]. Firstly, anatomical and physiological distinctions exist between the human spleen and murine counterparts. Secondly, the inherent risk of potentially severe intraperitoneal hemorrhage precludes invasive exploration of the human spleen through techniques such as biopsy or needle aspiration. Therefore, current research on the spleen and IES filtering function mainly includes three methods: *ex vivo* [15, 16, 17], *in vitro* [18, 19, 20, 21], and *in silico* [22, 23, 24]. Safeukui *et al*. investigated the spleen’s filtration function through *ex vivo* perfusion experiments involving normal human spleens and RBCs with varying degrees of surface area loss. Their findings demonstrated that treatment with increasing concentrations of lysophosphatidylcholine (LPC) resulted in dose-dependent reductions in RBC surface area, increased osmotic fragility, and decreased deformability[15]. Furthermore, several research groups have developed spleen-on-chip platforms for *in vitro* studies to understand the spleen’s homeostatic balance [19, 18]. Additionally, a few numerical methods, including dissipative particle dynamics (DPD) [22, 24, 25, 26], smoothed dissipative particle dynamics (SDPD) [27], and boundary integral method [28, 29], have been employed to investigate the passage of RBCs through spleen-like slits.

While significant research has been dedicated to understanding the filtration process of RBCs by the spleen [15, 16, 22, 29, 24, 26], current investigations encounter two primary limitations. Firstly, existing *in vitro* and *in vivo* studies often treat endothelial cells as non-deformable when examining individual IES [25, 24], leading to a higher pressure differential required for cell traversal through these rigid slits [23]. This elevated pressure difference can potentially result in cellular damage as cells navigate through narrower slits. Previous research has evaluated the stiffness of various endothelial cells, revealing that endothelial cells are more prone to deformation under blood pressure [30, 31, 32]. Consequently, as RBCs pass through the slits, endothelial cells undergo some level of deformation, mitigating excessive cellular injury. Secondly, the precise filtration mechanism of the spleen sinus for cell suspensions remains insufficiently elucidated. Considering the human spleen’s high hematocrit (HCT) of approximately 80% and the intricate spleen environment [33], assessing the spleen using a singular filtration mechanism may not be sufficient. While the roles of ATP, pressure, and shear force in spleen filtration function are recognized [34, 35], contemporary research, whether conducted *in vitro* or *in silico*, predominantly relies on passive mechanisms where cell suspensions are passively compressed through the slits. However, prior studies suggested that the endothelial cells lining the spleen sinus can intermittently open due to various factors, promoting RBC passage and potentially preventing splenic congestion.

This research employs a cellular-scale model of RBCs and utilizes the DPD method to examine the filtration mechanism of IES by considering its deformability. Initially, the worm-like string model is used to characterize endothelial cells, aiming to elucidate the distinctions between deformable and rigid models in single-cell filtration dynamics through IES. Furthermore, various hypotheses are integrated to construct an active model for the filtration of cell suspensions. Within this active model, the filtration dynamics of cell suspensions is investigated, and a comparative analysis is conducted among active, passive, and rigid models. This study also pioneers the quantification of IES deformation and employs a more precise model to investigate the RBC dynamics within the spleen. Additionally, the impact of cell surface area-volume ratio on filtration function is evaluated, and the effects of spherical shape and stiffness on filtration are quantified, with an additional consideration of their relevance to malaria disease. The proposed model offers insights for disease analysis and prediction, contributing to potential preventive strategies.

The structure of this paper is outlined as follows. Section 2 provides an overview of the models and methods utilized, encompassing the problem description (Section 2.1), the DPD method and multiscale blood cell models (Section 2.2), the passive deformable IES model (Section 2.3), and the active deformable IES model (Section 2.4). Section 3 presents the results, commencing with two validation studies (Section 3.1) focused on the calibration of endothelial cell stiffness and the determination of the critical shear modulus governing RBC passage through the IES. Subsequently, Section 3.2 examines the influence of RBC surface area to volume ratio on their retention by deformable IES, while Section 3.3 delves into the impact of RBC sphericity, stiffness, and crossing-slit pressure on retention. Section 3.4 investigates the dynamics of malaria-infected RBCs traversing deformable IES, followed by the quantification of RBC suspension dynamics through an array of slits in Section 3.5. The simulation of the filtration function of the 3D splenon is discussed in Section 3.6. Finally, Section 4 encompasses the conclusion and discussion of the findings.

## 2. Methods and models

### 2.1. Problem description

The left panel of Figure 1 illustrates the three-dimensional structure of the spleen sinus, demonstrating the migration of cells from the spleen cords to the spleen lumen within IES. This includes endothelial cells, fibers, and erythrocytes. The dimensions and geometry of the spleen sinus fall within the reported range [13]. The right section examines the spleen sinus in both *in vivo* and *in silico* conditions. Given the well-known caliber of approximately 1 *μ*m, the passage of cells through the slit poses a challenge. Consequently, when cells traverse the slit, there is observable deformation in the endothelial cells. This deformation proves advantageous for the cell’s passage through the slit. The right panel of Figure 1 visually captures the deformation of endothelial cells during cell passage through the slit.

**Figure 1:**
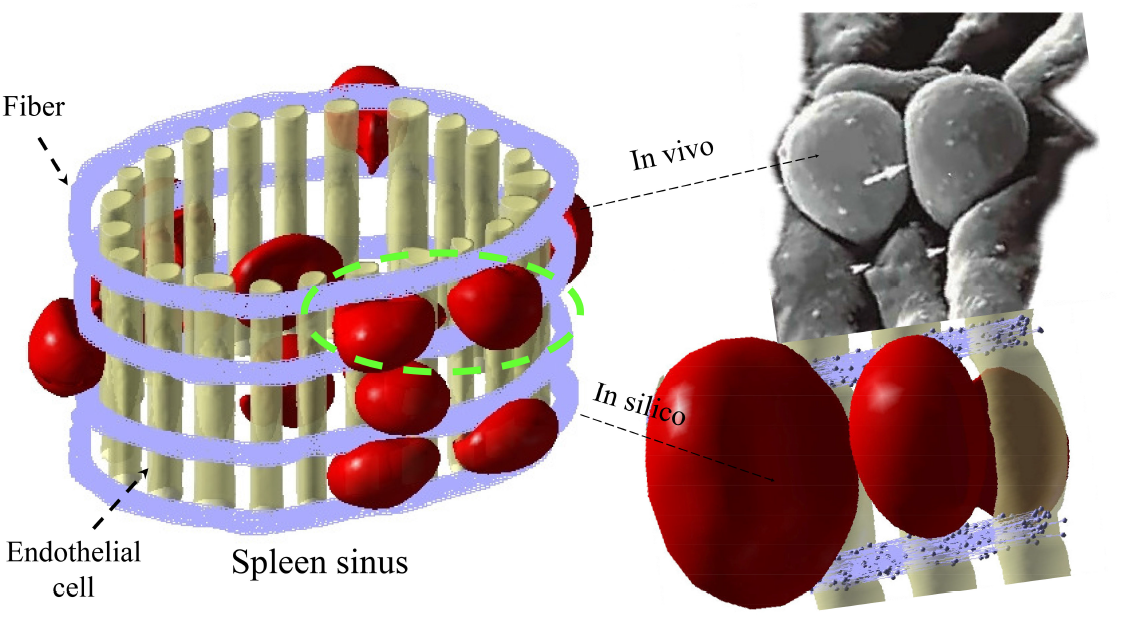
Simulation setup of the deformable 3D splenon. The left panel portrays a segment of a three-dimensional spleen sinus, comprising endothelial cells, fibers, and RBCs, wherein the RBCs traverse IES to transit from the cords to the spleen lumen. The right panel provides a comparison of an RBC squeezing through a slit *in vivo* and a simulation of the same process using a deformable IES model.

### 2.2. DPD method and multiscale blood cell models

In this study, we utilize DPD to simulate plasma, RBCs, and macrophages. DPD is a mesoscopic particle-based simulation technique, where each DPD particle represents an aggregate of molecules interacting with others through soft pairwise forces [36, 37]. This method accurately captures the hydrodynamic behavior of fluids at the mesoscale and has proven successful in investigating complex fluid systems [38, 21, 39, 40]. The evolution of computational capabilities in the past two decades has fostered the advancement and application of multiscale biophysical models for RBCs. This includes models operating at the protein level [41, 42, 43, 44, 45, 46, 47, 48, 49] and cellular level [**?** 44, 50, 51, 52, 53, 54, 55, 56, 57, 40, 58]. While protein-level RBC models can simulate pathological alterations in RBC membrane structure associated with blood disorders, their application to modeling blood cell suspensions or blood flow is hindered by computational costs. Building upon our prior research on sickle cell adhesion [59, 60, 20], we implement a cellular-level model [61] based on DPD [62] to simulate normal and sickle RBCs, as well as macrophages (for additional details, refer to section S1).

### 2.3. Passive deformable IES model

In prior research [22, 25], the endothelial cell has conventionally been treated as a rigid wall. However, it has been observed to undergo deformation in response to shear flow and in micropipette experiments [30, 31, 32]. Consequently, we employ a passive model to replicate the deformation of the spleen sinus during cell traversal through the slit. The configuration in Figure 2A depicts an RBC navigating IES, wherein we employ the worm-like string model for the endothelial cell, a model widely utilized in diverse works [63, 64, 65]. The endothelial cell’s surface is represented by a 2D triangulated network comprising *N*_*v*_ vertices (DPD particles). Elastic bonds connecting these vertices, totaling *N*_*s*_, are introduced to enforce appropriate membrane mechanics. The free energy (*V*_*cell*_) is expressed as follows:

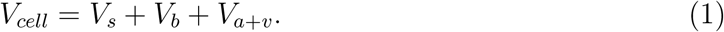

**Figure 2:**
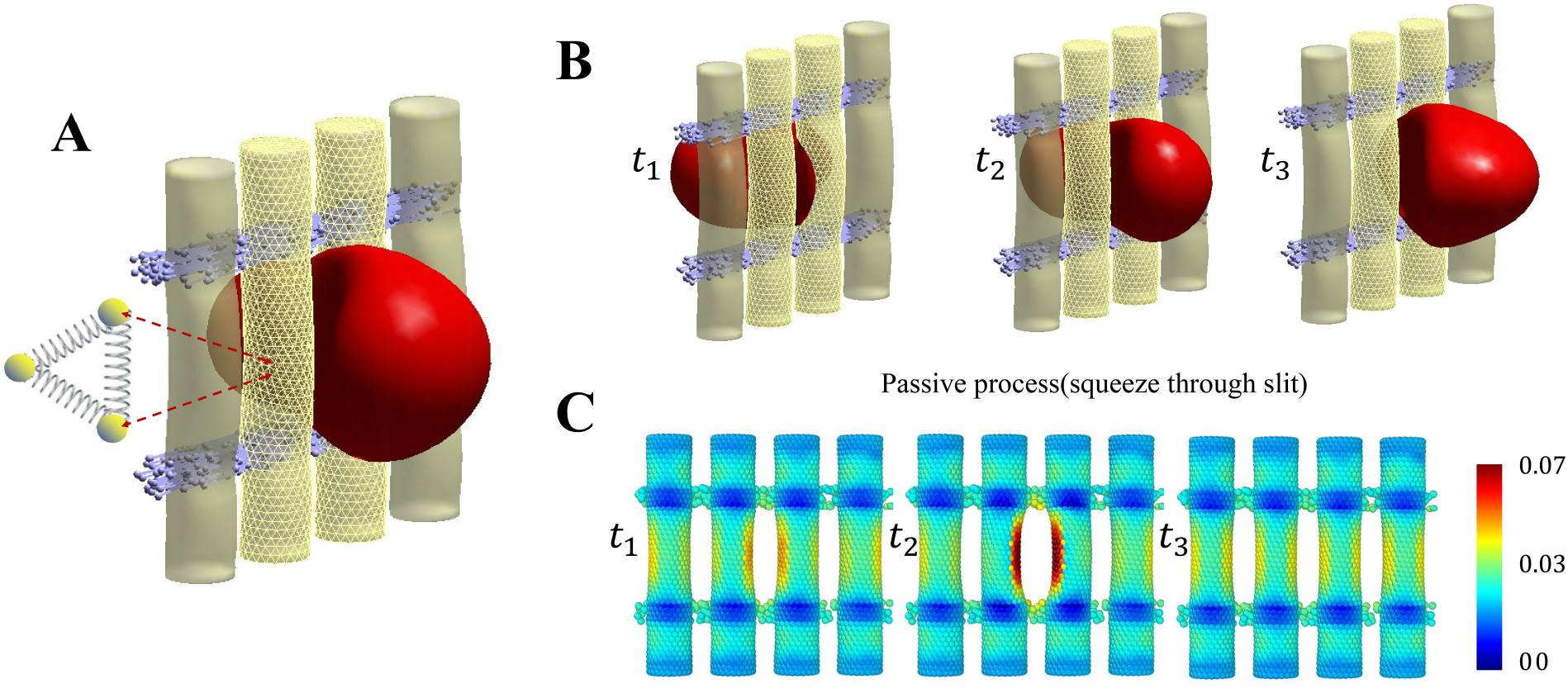
Passive deformable IES model where the deformation of IES is dictated by its interaction with traversing RBCs. (A) Illustration of the RBC navigating a deformable slit, with the endothelial cell’s fiber depicted as interconnected strings. (B) The temporal span *t*_1_ ∼ *t*_3_ encapsulates the sequential snapshots of an RBC traversing the passive IES model. (C) The magnitude of the volumetric strain of the endothelial cell measured in the three snapshots recorded in (B).

The elastic energy *V*_*s*_ representing the elastic interactions of the cell membrane is defined by

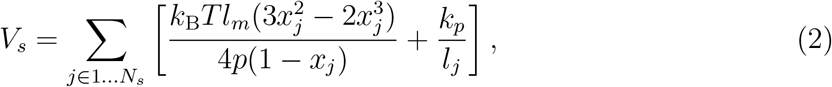

where *p* is the persistence length, *k*_*p*_ is the spring constant, *k*_B_*T* is the energy unit, *l*_*j*_ is the length of the spring *j, l*_*m*_ is the maximum spring extension, and *x*_*j*_ = *l*_*j*_*/l*_*m*_. *p* and *k*_*p*_ are computed by balancing the forces at equilibrium and from their relation to the macroscopic shear modulus, *μ*_*s*_:

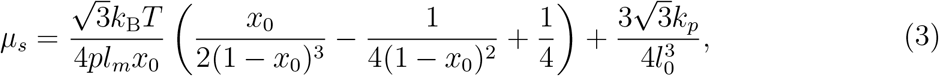

where *l*_0_ is the equilibrium spring length and *x*_0_ = *l*_0_*/l*_*m*_. The bending resistance *V*_*b*_ of the cell membrane is modeled by

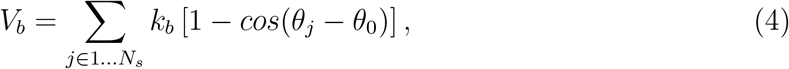

where *k*_*b*_ is the bending constant, and it is related to the macroscopic bending rigidity *k*_*c*_ with the expression 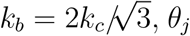 is the instantaneous angle between two adjacent triangles sharing the common cedge *j*, and *θ*_0_ is the spontaneous angle. In addition, the area and volume constraints *V*_*a*+*v*_ are imposed to mimic the area-preserving lipid bilayer and the incompressible interior fluid. The corresponding energy is given by

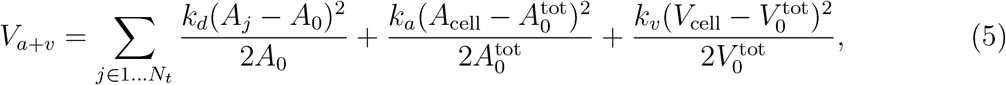

where *N*_*t*_ is the number of triangles in the membrane network, *A*_0_ is the equilibrium value of a triangle area, and *k*_*d*_, *k*_*a*_ and *k*_*v*_ are the local area, global area and volume constraint coefficients, respectively. The terms 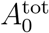 and 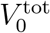 are targeted cell area and volume.

Figure 2B illustrates the temporal evolution spanning *t*_1_ to *t*_3_, encompassing distinct phases within the passive model. During this interval, the RBC undergoes deformation while traversing the compliant slit. Notably, the deformation of the endothelial cell is evident when the RBC remains within the slit. The alignment of the volumetric strain of the endothelial cell with the events described in Figure 2B is demonstrated in Figure 2C. A comprehensive account of the calculation process is provided in the supplementary materials. The analysis reveals that the tensor of the endothelial cell initially increases and subsequently returns to its initial state after the cell through out of the slit.

### 2.4. Active deformable IES model

In addition to the passive model, an active model manifests when the cell suspension traverses the sinus. The active model is underpinned by three hypotheses. Firstly, variations in venous slit caliber are governed by the prevailing “pressure” in the red pulp’s reticular meshwork at a specific moment [35]. Secondly, the slit caliber is influenced by the contractility of stress fibers in human endothelial cells, possibly triggered by ATP or shear force resulting from fluid flow [34]. Thirdly, there are no preformed apertures in the sinus walls; instead, slits between sinus endothelial cells, typically closed, widen as cells pass through them [8]. For our analysis, we consider only the first two hypotheses. Figure 3A illustrates the active model’s process in three stages: increasing upstream pressure, altering contractility, and actively opening slits. In contrast to the passive model, the active model exhibits two primary distinctions. Firstly, according to the pressure-dependent hypothesis, a pressure threshold for the upstream is established, and pressure is monitored at each timestep. If the pressure exceeds the designated threshold, the active process is initiated. Secondly, considering the second hypothesis, endothelial cell contractility undergoes modification. To realize this adjustment, the maximum spring extension *l*_*m*_ increases to 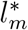, and the targeted cell area 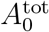 and volume 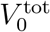 change to 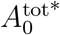 and 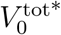, respectively. Consequently, the active endothelial cell alters contractility, thereby facilitating the passage of RBCs. The algorithm for the active process is summarized as follows.

**Figure 3:**
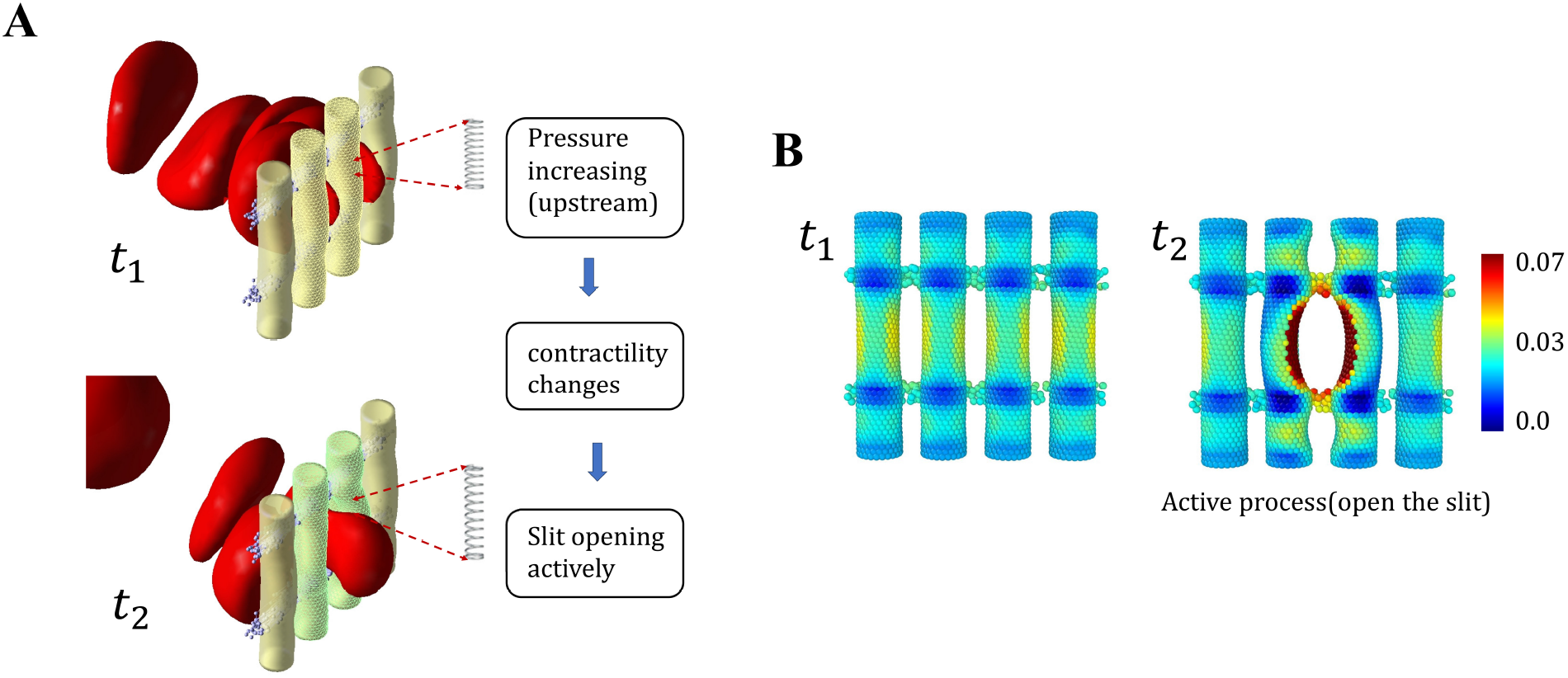
Active IES model where the deformation of IES is regulated by its surrounding pressure. (A) The temporal span *t*_1_ ∼ *t*_2_ illustrates that the active model functions through the following three steps: 1) upstream pressure elevation; 2) alterations in fiber contractility; 3) active opening of the slit. (B) The volumetric magnitude of strain tensor of the endothelial cell measured at snapshot *t*_2_ in (A).

Step 1: Update the pressure of the upstream.

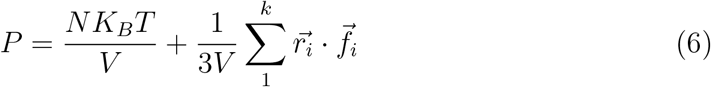

where *N* is the number of atoms in the system, *K*_*B*_ is the Boltzmann constant, *T* is the temperature, and V is the system volume. The second term is the virial, where 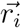 and 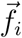 are the position and force vector of atom *i. k* necessarily includes atoms from neighboring subdomains (so-called ghost atoms), and the position and force vectors of ghost atoms are thus included in the summation.

Step 2: Examine if the upstream pressure exceeds the threshold(*P > P* ^*^), where *P* ^*^ is the threshold pressure.

Step 3: Activate the active process, switch to the active maximum spring extension parameter 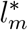 for elastic interaction energy,

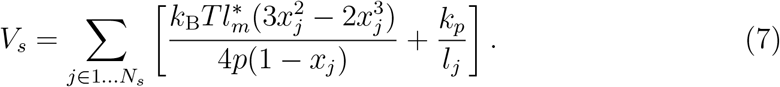

Step 4: Switch to the targeted area surface and volume 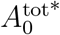 and 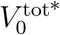.

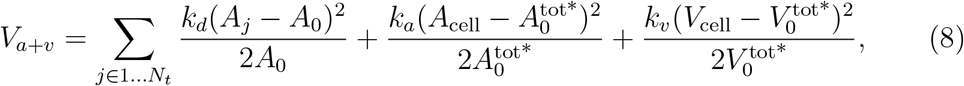

Figure 3B shows that the temporal span *t*_1_ ∼ *t*_3_ encapsulates the successive phases of the active model: *t*_1_ depicts the pressure increase due to the accumulation of the RBCs, the endothelial cell actively opens as the RBC traverses the deformable slit at *t*_2_, and the endothelial cell remains active after the cell passes through the slit at *t*_3_, the active process would appear to have durations of ∼ O(10s). Figure 3 shows the volumetric strain of the endothelial cell corresponds to the events outlined in B.

The extended code derived based on LAMMPS is employed to compute all simulations. Each simulation involves approximately 2,000,000 time steps and necessitates 960 CPU core hours. These computations utilize the computational resources available at the Center for Computation and Visualization at Brown University, featuring Intel Xeon E5-2670 2.6 GHz 24-core processors.

## 3. Results

### 3.1. Validation

#### 3.1.1. Calibration of the stiffness of endothelial cell

Numerous works have explored the stiffness of diverse endothelial cells, employing the micropipette-aspiration technique to determine the elastic modulus. Mohammadkarim *et al*. executed a micropipette-aspiration experiment to assess the stiffness of mechanical properties in human umbilical vein endothelial cells across control and radiation-induced samples [32]. In Figure 4A, the geometry and dimensions of the micropipette setup are illustrated, where the endothelial cell exhibits a diameter of around 10 *μm*, and the inner diameter of the micropipette measures 3 *μm*. Varied pressure drops, namely 103.67*Pa*, 179.33*Pa*, 236.07*Pa*, and 340.40*Pa* were applied to aspirate the endothelial cell, resulting in corresponding aspirated lengths of 0.89*μm*, 1.62*μm*, 2.06*μm*, and 2.8*μm*. In our simulation, we selected a set of shear modulus and bending modulus values, specifically *Es* = 7.010^−5^*N/m* and *Eb* = 8.710^−19^*J*, to calibrate parameters matching the experimental outcomes. This value is also consistent with other experimental results [30, 31]. Figure 4B displays snapshots from various simulations, while Figure 4C illustrates the aspiration ratio’s variation with pressure drops, showcasing the simulation results’ alignment with experimental findings.

**Figure 4:**
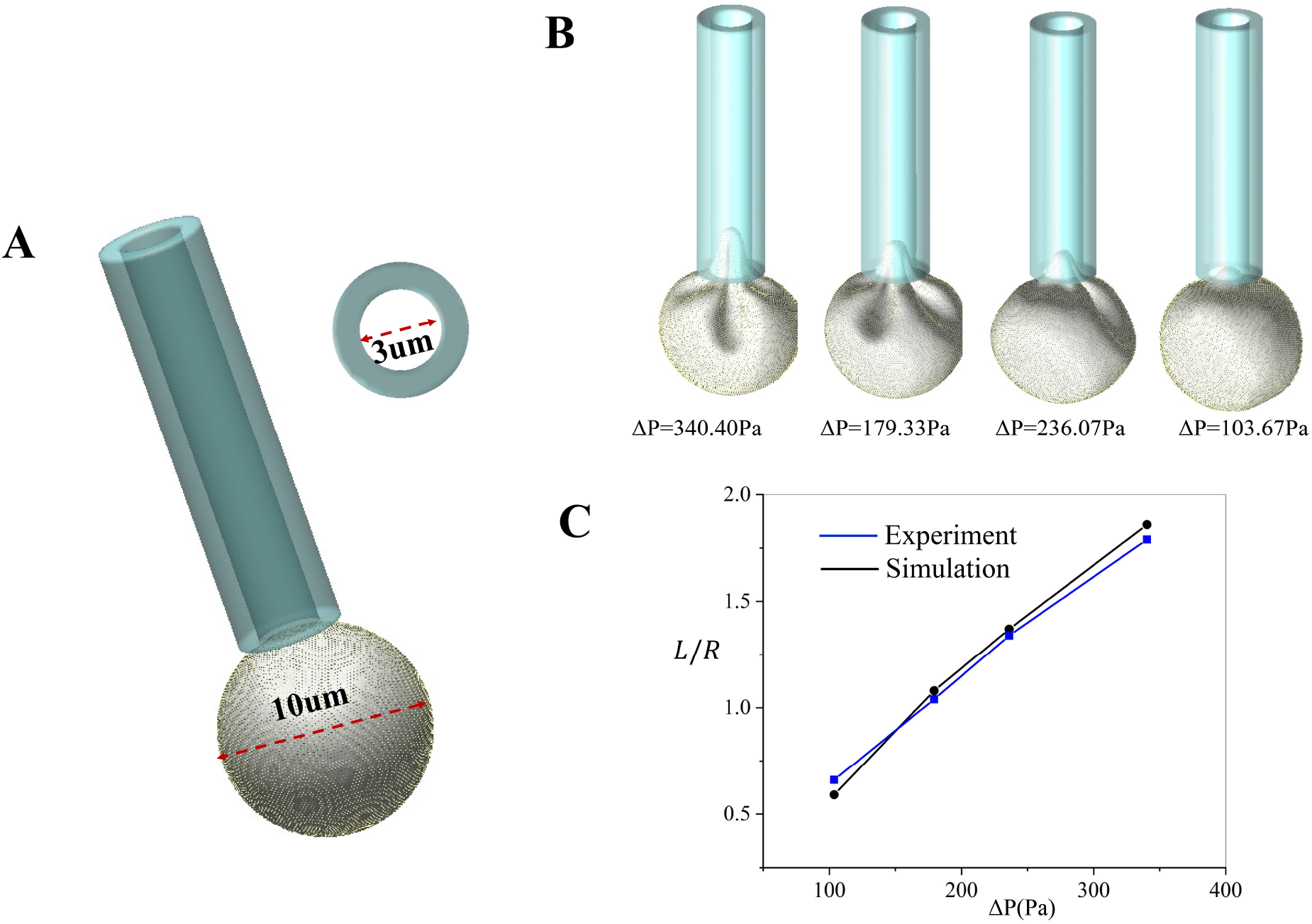
Calibration of the endothelium cell model using micropipette aspiration. (A) The geometry and dimensions of the micropipette and the endothelial cell. (B) Variations in aspiration length of the endothelial cell are observed under different pressure conditions. (C) Both experimental and simulation data demonstrate alterations in the aspiration ratio across a range of pressures [32]. The experiment from Mohammadkarim *et al*.

#### 3.1.2. Determination of the critical shear modulus dictating the RBC passage of the IES

Subsequently, we conducted a comprehensive analysis to ascertain the probability of an individual RBC either navigating through a micro-slit or undergoing retention while traversing diverse geometric configurations such as cylinders, squares, and elongated shapes. The dimensions and configurations of the slits are delineated in the right panel of Figure 5. A pressure gradient was also applied to induce an upstream fluid velocity of approximately 100*μm/s*. The critical shear modulus of the RBC varied within the range of 6.0 to 22.8 *μN/m*. The hierarchy among different geometries was established as follows: circular (d=1.5) *<* circular(d=1.8) *<* square (d=2) ≈ elongated *<* square(d=2.4). A comparative analysis with Qiang’s findings is depicted in the left panel of Figure 5, demonstrating consistency between our results and previous observations [19]. Moreover, the critical shear modulus required for RBCs to traverse through the elongated slit was estimated using the Young–Laplace equation 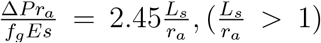, incorporating a geometric adjustment factor representing the minor radius of the RBC. Δ*P* is the constant pressure difference. *L*_*a*_ is the maximum value of its major axis of RBC in the slit. *E*_*s*_ is the estimated shear modulus, and *r*_*a*_ is the radius of the short axis of RBC. As the first order approximation, *f*_*g*_ = 1 is taken in this study for estimating the critical shear modulus [66]. The calculated critical shear modulus is 14.6*μN/m*, aligning closely with our obtained results.

**Figure 5:**
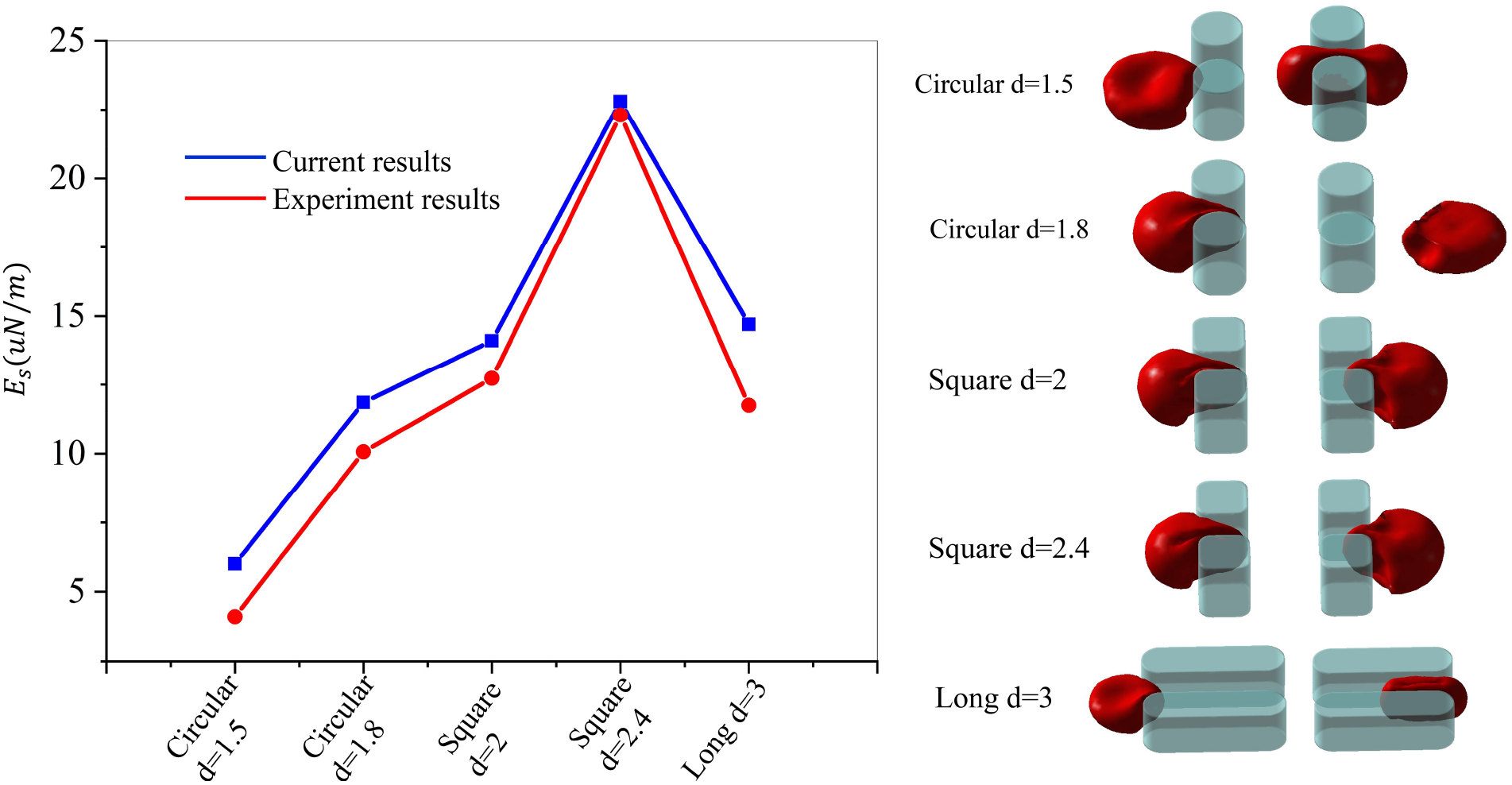
Impact of the size and shape of the rigid IES on critical shear modulus of RBCs dictating their IES retention. The left diagram illustrates the critical shear modulus of the RBCs measured from microfluidic experiments [19] and simulations above which RBCs will be retained by IES. The snapshots on the right capture instances of RBC entry and exit slits with diverse geometries. The experiment from Qiang *et al*.

### 3.2. Explore the impact of surface area to volume ratio of RBCs on their retention by deformable IES

The range of size distribution observed in RBCs within the context of normal human subjects is notably diverse, encompassing cell surface areas ranging from 80 to 180 *μm*^2^, and volumes varying between 60 and 160 *μm*^3^ [67, 68]. Microsphere experiments and studies involving isolated perfused human spleens have estimated a pressure gradient of 1.0*Pa/μm* sufficient to facilitate the passage of RBCs of varying sizes through the spleen sinus. To align our analysis with experimental findings, we conducted simulations employing both rigid and passive models to examine the transit of normal RBCs with different sizes through the IES under a fixed pressure gradient of 1.0*Pa/μm* [15, 69]. Figure 6A illustrates the dimensions and configuration of the equipment, while representative shape transitions of an RBC passing through the rigid and passive models are presented in Figure 6B and C. We observed that cells undergo deformation when squeezed through both models. In comparison with the rigid model, endothelial cells also undergo slight deformation due to the squeezing action of RBCs in the passive model at *t* = 85 and 130*ms*, thereby increasing the width of the slits, facilitating cell passage. Consequently, the time required for RBCs to pass through the passive model is shorter than that required to pass through the rigid model.

**Figure 6:**
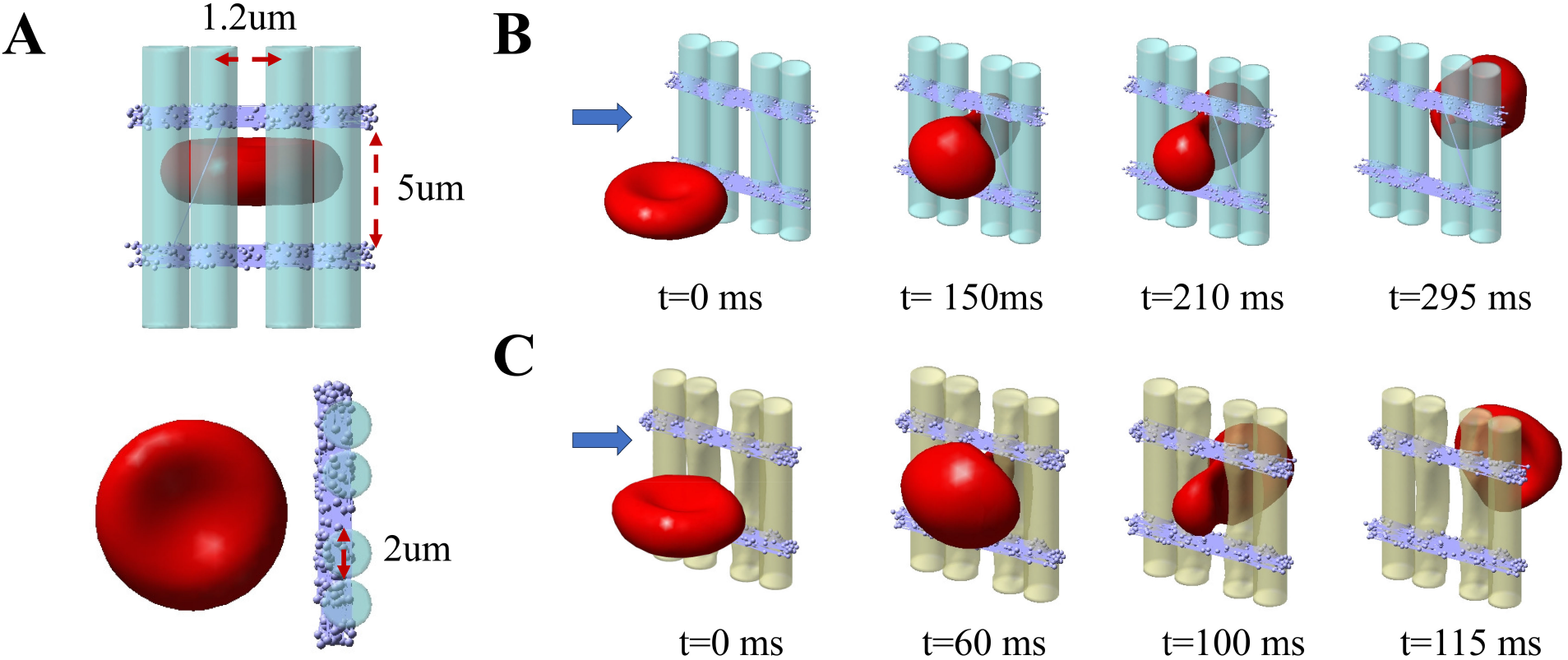
Computational simulation depicting the passage of an RBC through deformable IES. (A) Schematic representation of the geometry and dimensions of the splenic IES. (B) Sequential snapshots illustrating the time interval of RBC traversal under the passive model. (C) Time interval snapshots portraying RBC passage under the active model.

We have identified the threshold volume for efficient traversal of RBCs based on their surface area, demonstrating that cells below this critical volume can effectively traverse the IES, whereas larger cells encounter impediments during crossing. Initially, we applied a theoretical model elucidated by Canham and Burton previously to estimate the critical minimum surface area for a fixed cell volume[67]. Within the theoretical framework, the indispensable geometric prerequisites for an RBC to navigate through a slit of arbitrary width, length, and depth necessitate the cell to have adequate surface area enabling the formation of two spheres on both sides of the slit aperture, linked by an “infinitely” thin tether. The arrangement characterized by the maximum sphericity index, facilitating the passage of RBCs through a slit regardless of its dimensions, is achieved when the cell is evenly divided into two spheres positioned on either side of the slit. A straightforward calculation of the relationship between surface area (*S*) and the volume (*V*) of two equal spheres yields *S* = (72*πV* ^2^)^1*/*3^ = 6.09*V* ^2*/*3^ with a sphericity index (SI) of 0.7937. To adjust for the physiological conditions of RBCs at 37°C in the spleen, we initially increased their surface area by 2%, representing the surface dilation of the cell membrane at 37°C [70]. Subsequently, we increased it by an additional 2% to account for the cell membrane’s ability to expand slightly under tension without breaking [71]. Notably, almost all cells exhibit, for a given volume, a surface area at least equal to or higher than that required to form two equal spheres and a tether.

The black solid line represents the theoretical prediction depicted in Figure 7A and B, the red dashed line illustrates the simulation outcomes of the rigid model in Figure 7A, and the blue dashed line in Figure 7B corresponds to the results of the passive model. In this context, RBCs characterized by surface area and volume coordinates situated to the right of this solid line could not traverse the IES, and the RBCs positioned to the left side can be circulated in the bloodstream. Notably, the passive model demonstrates better alignment with the theoretical prediction than the rigid model. Furthermore, the sphericity of RBCs plays a pivotal role in the spleen’s filtration function. The simulation results align with more recent data obtained using a high-throughput device consisting of thousands of parallel microchannels to measure RBC surface area and volume [68]. These results indicate that nearly all RBCs can traverse the interendothelial slits in the spleen. However, a subset of RBCs, characterized by smaller volume and surface area, faces obstruction by the interendothelial slits; these cells are likely to be older and more rigid. Alternatively, these senescent cells may represent a minor fraction that has just reached the physical limits triggering retention but has not yet been directed to the filtering beds of the spleen.

**Figure 7:**
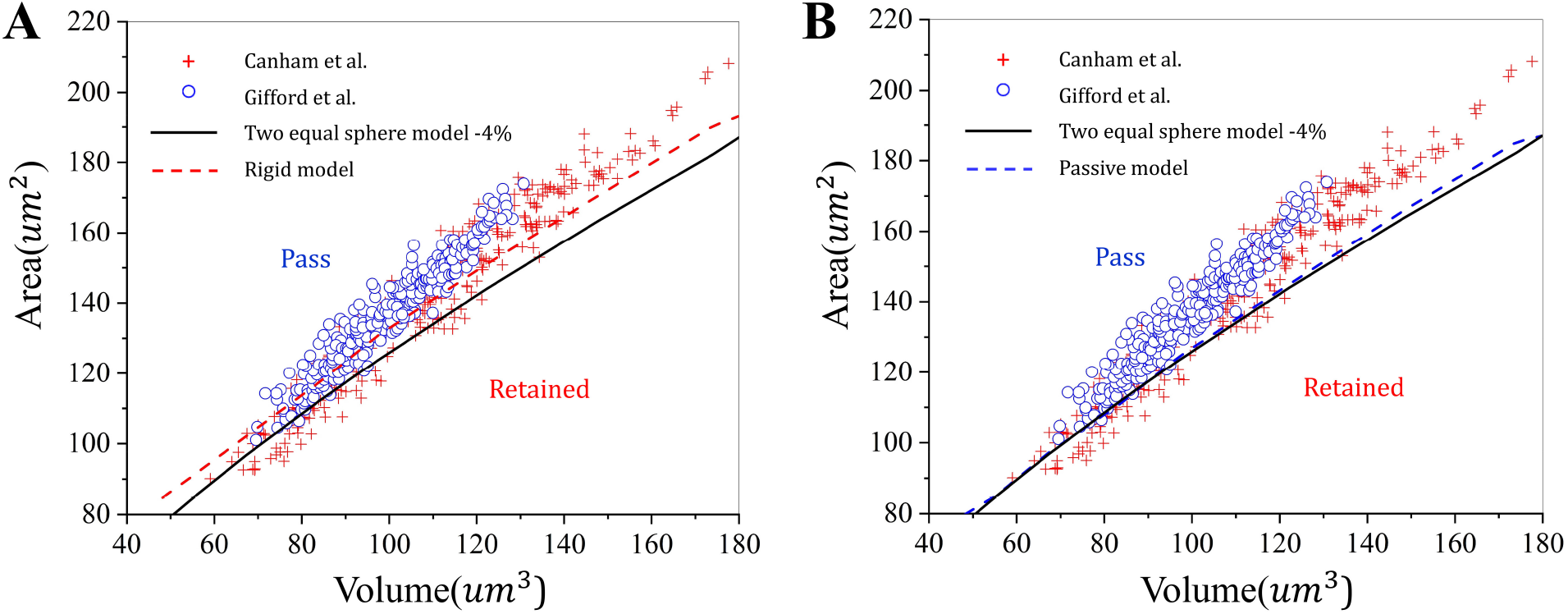
Projected correlation between the volume and surface area of normal RBCs under a pressure gradient of 1*Pa/μm*. (A) The minimum surface area required for RBCs with varying volumes transiting through rigid IES. (B) The minimum surface area required for RBCs with varying volume transiting through deformable IES. The scatter symbols are adopted from the work of Canham et al. [67] and Gifford et al. [68].

### 3.3. Examine the effect of sphericity and Stiffness of RBCs and the crossing-slit pressure on RBC retention by deformable IES

Over the approximately 120-day lifespan of RBCs, they gradually accumulate damage. Towards the end of their life cycle, RBCs undergo detrimental changes, causing a loss of deformability that hinders their ability to pass through the IES in the red pulp [16, 72, 12]. Various independent *in vitro*[13, 18, 73, 74] and *ex vivo* [75, 15] experimental studies have demonstrated that RBCs exhibiting increased stiffness and sphericity are more prone to mechanical retention at IES. The spleen cord environments are complex, with varying pressure distributions. Therefore, we aim to investigate the influence of stiffness and sphericity on filtration mechanics. Figure 8A illustrates the minimum pressure gradient required for RBC passage through the slit relative to RBC sphericity. Notably, lower pressure gradients are observed for the passive model than the rigid model. Subsequently, pressure gradients increase with sphericity, with a more pronounced dependence when sphericity exceeds 0.76 for both rigid and passive models. Recently, Alexis *et al*. clarified the mechanisms influencing the dynamics of RBC passage or retention within a narrow slit through the use of multiscale modeling, and their findings also indicate a noticeable increase in the minimum pressure gradient when the sphericity of RBCs exceeds 0.76 [23]. Figure 8B depicts changes in the minimum pressure gradient over RBC stiffness for both rigid and passive models. The results indicate an increase in the minimum pressure gradient with a higher shear modulus. However, the plot reaches a plateau when the shear modulus exceeds five times that of a normal cell. Compared to sphericity, stiffness has a lesser impact on the filtration process.

**Figure 8:**
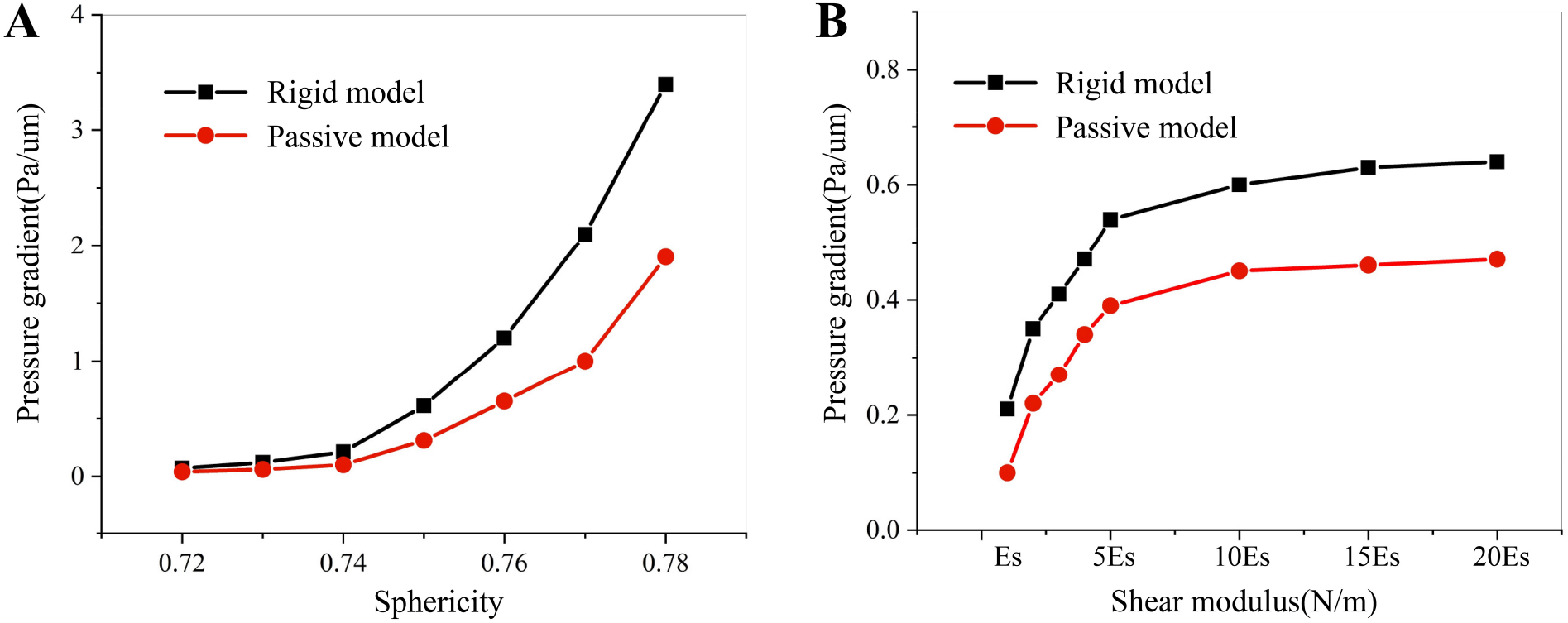
The impact of RBC sphericity and shear modulus on the critical pressure driving them through IES. (A) The critical pressure gradient for driving normal cells with varying sphericity through IES. (B) The critical pressure gradient for driving normal cells with varying shear modulus through IES. The black line denotes the rigid model, while the red line represents the passive model.

### 3.4. Investigate the dynamics of malaria-infected RBCs traversing through deformable IES

To quantify the transit time when RBC traverses the slit in the rigid and passive models, we design a microfluidic chip, which is shown in Figure 9A. We now consider normal, ring, and trophozoite-stage RBCs invaded by P. falciparum merozoites; the surface area of the RBCs in these three cases, respectively, was taken to be 135, 122.04, and 115.83*μm*^2^, and the respective values of the membrane shear modulus were 5.5, 15.5, and 28.9 *pN/μm*. The cell volume was taken to be 94*μm*^3^ for all three cases. Figure 9B shows the velocity changes over time for the normal cell flow through the slit in the microfluidic chip. When the applied velocity at the upstream is about 100 *μm/s*, the transit time in the passive model is 110 ms; however, it will take 310 ms in the rigid model. We also estimate the transit time of different stage RBCs invaded by P. falciparum merozoite in Figure 9C. We can find that it is easier for the RBCs invaded by P. falciparum merozoites to transit the slit in the passive model. The ring stage cell cannot pass through the slit in the rigid model, but it can pass through the passive model. During the trophozoite-stage RBCs, the cell cannot pass through the slit both in rigid and passive models. We also checked the stiffness of endothelial cell influence on the transit time for the normal cell, shown in Figure 9D. The phase diagram shows the transit time changes with the shear and bending modulus. We can find that the transit time will increase with the shear and bending modulus. When we increase the shear and bending modulus to 100 times normal RBCs, the transit time will be about three times when the stiffness is the same as that of a normal cell.

**Figure 9:**
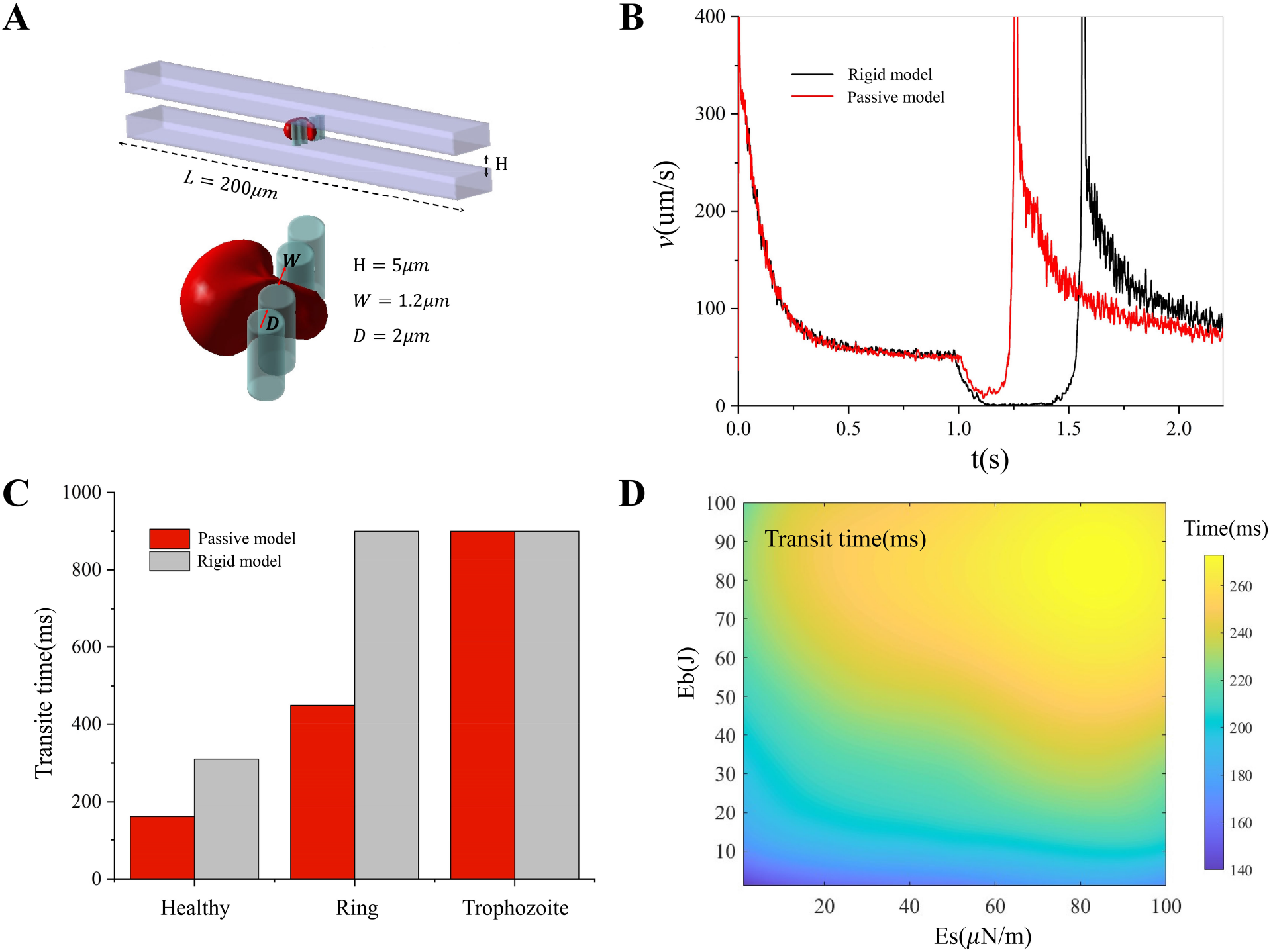
Simulation of an RBC traversing a single slit. (A) Simulation setup for modeling an RBC passing through a single slit. (B) The temporal variation in the velocity of a normal RBC as it traverses rigid and deformable slits. (C) Evaluating the transit time of P. falciparum-infected RBCs at various stages passing through rigid and deformable slits. (D) Systematic investigation of the impact of shear and bending modulus of the endothelium cell on the transit time of normal RBCs.

### 3.5. Quantify the dynamics of RBC suspension traversing an array of slits

MacDonald *et al*. have approximated that hematocrit levels within the human spleen can reach up to 78% [33], with recent empirical findings suggesting notable heterogeneity in the size, shape, and mechanical characteristics of individual RBCs within the same population [68]. These variations carry potential implications for the overall health and biological functionality of the entire cell population. Consequently, we employed a cell suspension within the microfluidic chip to discern differences among the rigid, passive, and active models. Figure 10A depicts the dimensions and size of the microfluidic chip, with slits randomly distributed. The slit dimensions range between 0.5 ∼ 1.2*μm*, consistent with the reported data [13]. Illustrated in Figure 10B is the *in vivo* passage of the cell suspension through IES, indicating cellular accumulation in spleen cords, with some RBCs traversing the IES into the spleen lumen. In Figure 10C, the passage of the cell suspension through the slit is examined across three models, rigid model, passive model, and active model. Under consistent conditions, differences in the level of upstream cell blockage are evident. Specifically, there is a greater degree of cell blockage in rigid models than in passive and active models, wherein the extent of cell blockage is notably diminished. To accurately measure the degree of cell blockage, we performed a statistical analysis on the number of cells traversing the slits across time in Figure 10D. Our findings reveal that the number of cells passing through slits exhibits an oscillatory behavior across all three models. The phenomenon of cell passage through slits exhibiting oscillations has been previously observed both *in vivo* and *in vitro* [35, 19]. MacDonald *et al*. conducted a statistical analysis of the number of cells passing through the spleen over time and found non-uniform variations in cell passage through the spleen sinus [35]. Recently, Qiang *et al*. observed a similar phenomenon while filtering cell suspensions in a microfluidic chip [19]. There exists variability in the amplitude of these oscillations, with the passive model showing the lowest amplitude, followed by the rigid model, and finally, the active model. The oscillatory nature of the rigid model suggests that cell suspension through slits is primarily driven by the accumulation of RBCs, and cell passage through slits occurs only when a certain critical pressure threshold is reached, indicating that the pressure gradient at the slits is not continuous. The diminished oscillations observed in the passive model compared to the rigid model can be attributed to two primary factors. Firstly, the prevalent narrow slit, typically around 1.2 micrometers in width in this work, which represent the widest slit observed in vivo, presents minimal hindrance for cellular passage through these openings. Secondly, as delineated in Figure 8, the minimal pressure gradient requisite for cellular transit through the passive model is lower than that of the rigid model. Consequently, cells in the passive model can traverse the slit without necessitating significant cellular accumulation, thereby mitigating pronounced fluctuations in the number of cells passing through over time. However, a reduction in the aperture size would similarly induce noticeable oscillations in the passive model. The active model plays a crucial role in establishing the oscillatory pattern *in vivo* and offers advantages over the other two models regarding oscillation patterns. In summary, cell passage through constrictions *in vivo* is primarily governed by a combination of passive and active mechanisms, contributing to the optimal functioning of the spleen.

**Figure 10:**
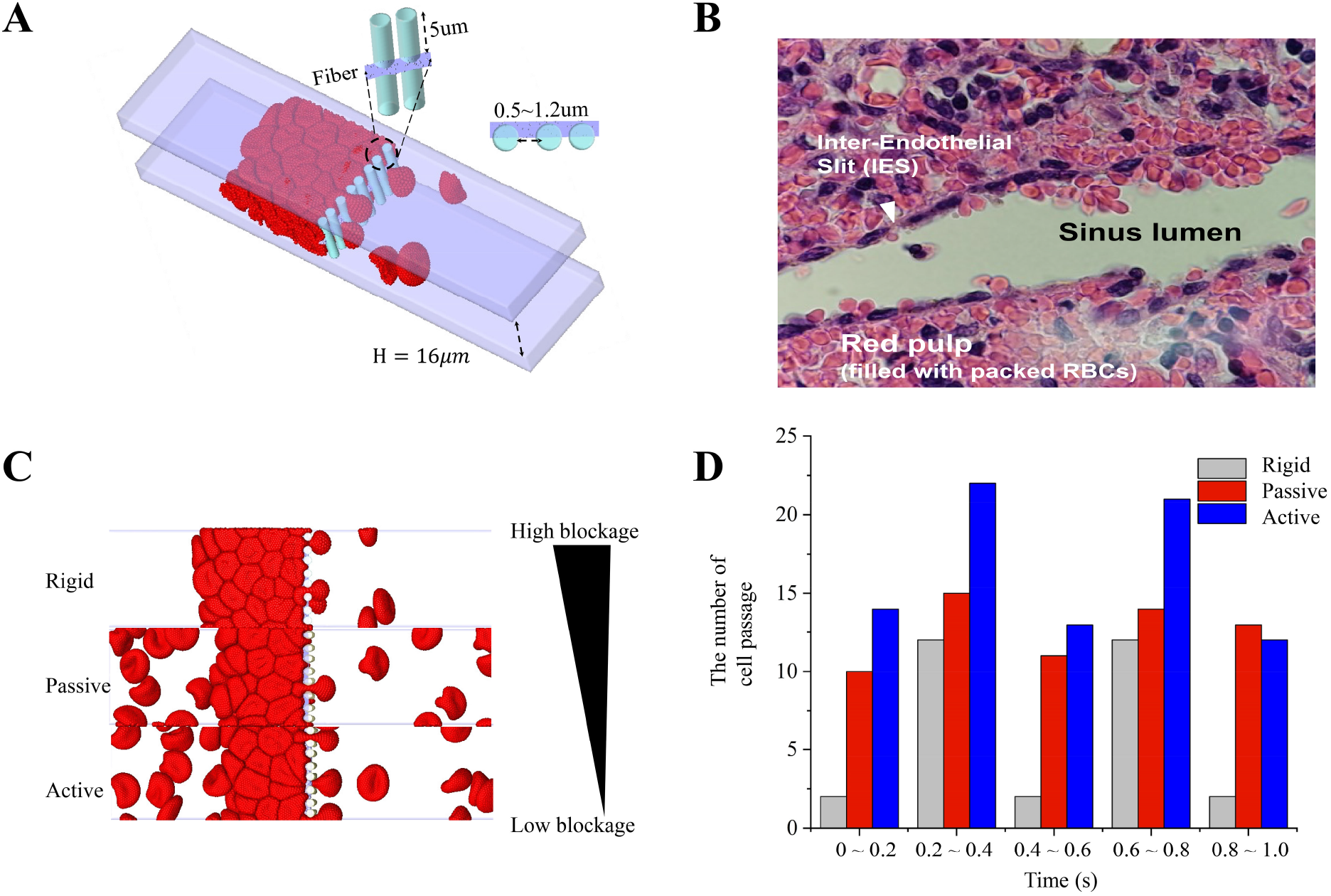
Simulation of RBC suspension traversing the microfluidic devices. (A) Illustration of the geometry and dimensions of the computational model of a microfluidic chip devised to mimic IES. (B) An *in vivo* image captures RBC suspension passage from spleen cords to the spleen lumen. reference. (C) Simulations of RBC suspension passing through microfluidic chips employing rigid, passive, and active models, highlighting varying degrees of blockage observed in the simulation results. (D) The statistical count of RBC traversing the slits changes over a simulation time of 1 second across three distinct slit models.

### 3.6. Simulation of the filtration function of the 3D splenon

To understand how the human spleen handles RBCs displaying decreased surface area at constant mean cell volume, isolated human spleens were perfused with a preparation containing RBCs exposed to different lysophosphatidylcholine (LPC) concentrations (1-2.5*μ*M, 3.5-5*μ*M, 6.0-7.5*μ*M, 8.5-15.0*μ*M,) and control RBCs [15], the surface area of the RBC will decrease with the increasing concentrations of LPC, but the volume of the RBC remains constant. The area and volume of the RBC under different concentrations of LPC are shown in Table 11. We keep the sphericity of RBC identical with the *ex vivo*. Figure 11A illustrates the simulation results of cells with varying degrees of sphericity within a three-dimensional spleen sinus. We observe that as the degree of cell sphericity increases, more cells are filtered in the spleen cords. Figure 11B depicts the retention rate of cells as a function of their sphericity. The level of splenic retention in the simulation(range, 18%-96%) and *ex vivo* (range, 19%-92%)increased with increasing extents of surface area loss. There was a positive correlation between the percentage of RBCs retained by the spleen and the extent of RBC surface area loss. A surface area loss of 8% resulted in the retention of 20% of RBCs within the spleen. Clearance of the RBC population became massive 96% after a surface area loss of 19%.

**Table 1:** Characteristics of treated cells with increasing concentrations of LPC in the simulation.

| Parameters | normal cell | 1 ∼ 2.5μM | 3.5 ∼ 5μM | 6.0 ∼ 7.5μM | 8.5 ∼ 15.0μM |
| --- | --- | --- | --- | --- | --- |
| Area(μm<sup>2</sup>) | 132.86 | 121.77 | 114.92 | 110.69 | 106.75 |
| Volume(μm<sup>3</sup>) | 92 | 92.66 | 94.21 | 97.73 | 96.33 |

**Figure 11:**
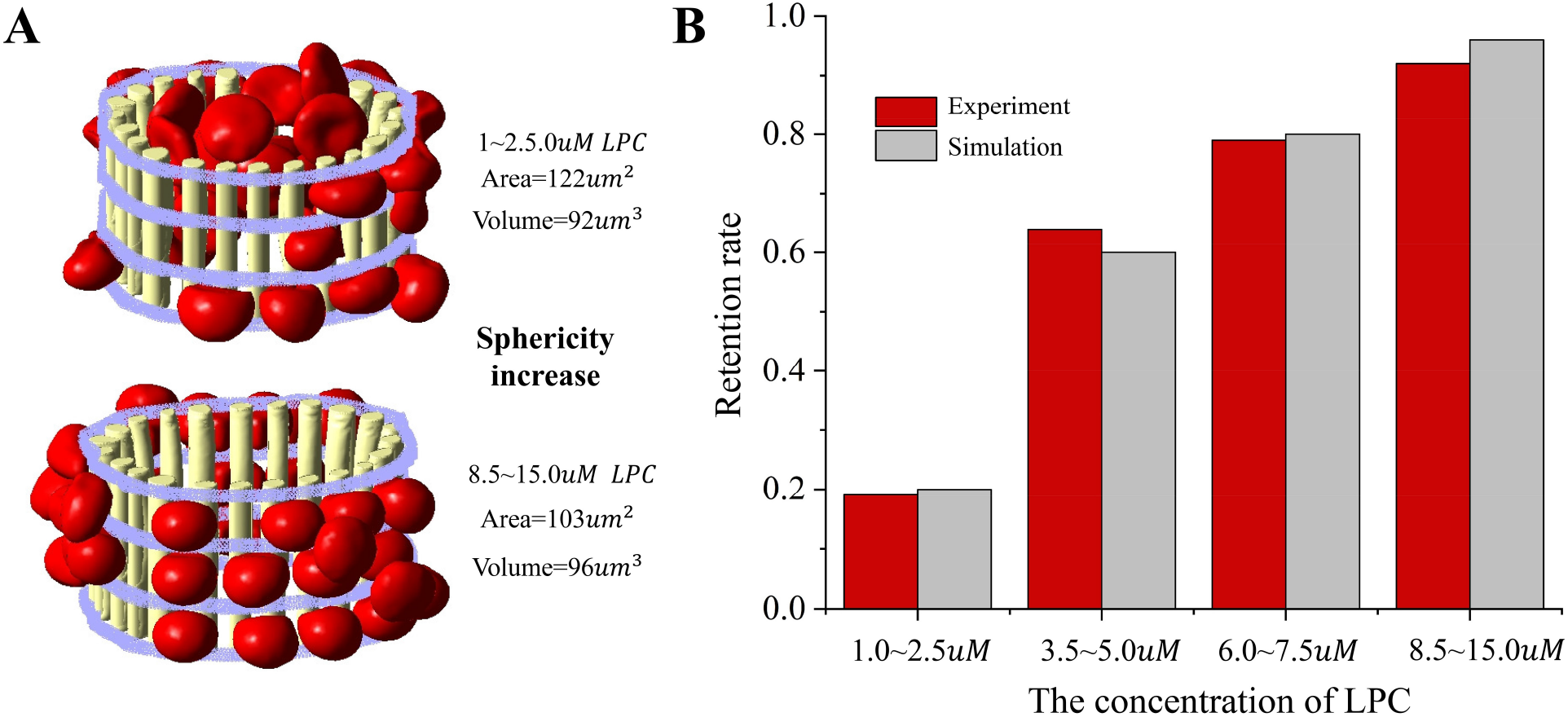
Simulation of the filtration function of the splenon. The mechanics of RBC retention are examined under *ex vivo* and *in silico* conditions, with increasing concentrations of LPC. (A) Simulation snapshots depict the traversal of RBC suspension through the 3D spleen sinus under varying RBC sphericity. (B) The comparison between simulation and *ex vivo* experimental results demonstrates the validity of the 3D splenon model.

## 4. Discussion and Summary

In this study, we initially propose two models, namely the passive model and the active model. In contrast to conventional rigid model [22, 25, 23], the passive model applies the worm-like string model to endothelial cells forming constrictions. Consequently, the endothelial cells deform due to their interaction with passing RBCs, thereby facilitating RBC transit. Although intensive research work has focused on studying individual RBCs passing through IES using *in vitro* and *in silico* approaches, the effort has typically been focusing on investigating non-deformable endothelial cells, which deviates from the actual *in vivo* environment [30, 31, 32]. The minimum pressure gradient required for cells to pass through IES exhibits an exponential growth pattern as the slit width decreases. This implies that in non-deformable models, cells would require a significantly large pressure difference to pass through IES when the slit width is below a certain threshold, posing a risk of cell rupture [23]. Therefore, there is an imminent need for a model that better reflects the mechanical characteristics of RBCs in the spleen environment to study and analyze cell mechanics in the spleen. In contrast to the passive model, the active model relies predominantly on the accumulation of upstream cells. This results in heightened pressure, prompting the contraction of the surface fibers of endothelial cells to create open channels. In the active model, cells require less energy to pass through IES, and the cell flux is boosted, effectively suppressing the possibility of the occurrence of splenomegaly. Unlike the passive model, cells from the active model do not need to squeeze through the slits with large deformation, which may result in the filtration failure of some aging and pathological cells. Therefore, it is likely that the primary mode of cell passage through IES in the human body remains passive [76], with the active mode serving as a supplementary mechanism to decongest the red pulp when it is crowded with diseased RBCs. The collaboration between active and passive models allows for the effective functioning of the spleen’s filtration capabilities. The passive model is predominantly involved in filtering aging and pathological cells, while the active model serves to alleviate pressure in the spleen.

Subsequently, we proceeded with the validation and calibration of the new models. Initially, we validated the passage of individual normal cells through the slit of different geometry. This was achieved by quantifying the minimum pressure gradient required for cell passage through the slit, thereby assessing the extent of cell passage through different slits. Our current work exhibits significant robustness compared to previous studies [19]. This validation serves as a reference for our new model, enabling a comprehensive quantification of the differences and advantages between the rigid and deformable models. Numerous studies have measured the stiffness of various endothelial cells under different conditions [30, 31, 32]. Due to the relatively low shear rate in the spleen, we chose to calibrate the stiffness of endothelial cells under control conditions [32]. This calibration resulted in the shear modulus and bending modulus of endothelial cells in the model as *Es* = 7.0 *×* 10^−5^*N/m* and *Eb* = 8.7 *×* 10^−19^*J*. Consequently, the mechanical properties of cells when squeezing through the slit differ significantly from previous studies [22, 25], as deformed endothelial cells facilitate cell passage. Finally, our focus shifted to cell suspensions, specifically examining the retention rates of cell suspensions with varying degrees of cell sphericity as they traverse the spleen. The cell retention rate increases with an increase in cell sphericity, consistent with previous experimental findings [15].

RBCs experience a circulation period of approximately 120 days in the human body, undergoing gradual aging and apoptosis [4]. Consequently, during this timeframe, RBCs exhibit diverse morphology and size distributions. Understanding the filtration function of the spleen requires accurate predictions of individual cell filtration characteristics through IES [3**?**]. In this study, we quantify the critical surface area for cell passage through the slit under varying volumes based on morphology. Under specific pressure conditions, most RBCs can pass through the spleen and enter the blood circulation, with only a few cells with small volumes and surface areas being filtered dynamically. The primary constituents of these cells are aging and apoptotic cells. Our simulation results of the deformable IES model closely align with theoretical predictions [67], and any disparities can be attributed to the oversight of surface area and volume considerations in the theoretical model. Consequently, the minimum surface area required for theoretical predictions is smaller than the outcomes obtained from DPD simulations. Earlier investigations by Pivkin *et al*. and Qi *et al*. explored the mechanics of single-cell filtration using a rigid model [22, 25], featuring a slit size and geometry similar to our present study. In contrast to their findings, the incorporation of a deformable slit in our model allows a broader spectrum of cells to pass through. In a more recent study, Peng’s research team examined the filtration mechanics of individual cells within a narrower IES [23], with the width approaching approximately 0.25*μm* in the experimental setup. In rigid models, normal cells require a minimum pressure of 2000 Pa to traverse such slits, potentially leading to substantial cellular damage. Conversely, in our deformable model, the passage of normal RBCs through small IES can be achieved with physiologically relevant crossing-slit pressure differences.

We also explored how the mechanical properties of individual cells and their sphericity affect their retention. The minimum pressure gradient necessary for cells to traverse the slit, depicted in Figure 6A, varies under different degrees of cell sphericity. The minimum pressure gradient required for cells to pass through the slit demonstrates an exponential increase with rising sphericity, particularly when the cell sphericity surpasses 0.76. Similarly, under diverse cell stiffness conditions, the minimum pressure gradient exhibits a logarithmic growth trend with the escalation of cell stiffness. By comparing the influence of cell sphericity and stiffness on cell mechanical properties, cell sphericity shows a more pronounced effect on the cell retention rate. This finding aligns with a recent investigation by Moreau *et al*., who examined the minimum pressure necessary for a single cell to navigate through a narrow slit [23].

To quantify the transit time of RBCs passing through the rigid and passive IES models, we considered normal, ring, and trophozoite stage RBCs infected by P. falciparum merozoites. It is observed that in the slit with geometry as illustrated in the Figure 9A, the normal cell can pass through in both models. However, the duration of their passage through the rigid model is almost two times longer than the passive model. Macdonald *et al*. conducted a statistical analysis of the time it takes for normal cells to traverse the slit *in vivo* [35]. Our observation aligns consistently with the findings reported in the previous study. Cells in the ring and trophozoite stages exhibit an inability to traverse the rigid slit. Conversely, in the passive model, only cells in the trophozoite stage face hindrances in passing through. In contrast to earlier investigations, our simulations of the mechanical attributes of malaria-infected cells within the spleen, employing a model that closely mimics the *in vivo* environment, suggest that the deformable model is more advantageous for facilitating cell passage through IES, a potential self-defense mechanism of the spleen to avoid congestion in the red pulp.

Finally, we employed rigid, passive, and active models to analyze the dynamics of cell suspensions traversing IES. Our results revealed variations in the degree of blockage in cell suspensions among the three models, with the severity of blockage following: rigid model *>* passive model *>* active model. In the human spleen, the collaborative interaction between the active and passive models may enhance the spleen’s filtration function. When cells experience blockage in the spleen, leading to increased pressure, the active model comes into play, preventing the occurrence of splenomegaly. Meanwhile, the passive model is responsible for filtering out pathologically altered cells. Our model aligns with the *in vivo* observations made by Macdonald et al. [35], who found that the number of cells passing through IES is not uniform [35]. Instead, it exhibits an oscillatory pattern with respect to time, supporting the findings from our simulations.

This study is subject to certain limitations. Firstly, our active model relies on multiple assumptions and lacks sufficient *in vivo* data for thorough model validation. This limitation stems from the intricate environment within the human spleen, and a singular model may not adequately capture the filtration patterns in this organ. Consequently, combining multiple models is imperative for a more precise representation. Secondly, our emphasis has predominantly centered on elucidating the mechanical properties of cells in relatively wide slits, with no simulations conducted for cells in narrower slits. The reported width range of IES in the spleen is between 0.25 ∼ 1.2*μm*, and our selection of 1.2*μm* represents the widest slit. Further research is warranted to explore the mechanical properties of cells in narrower slits within this specified range.

## Author Contributions

G. L.: designed research, performed research, contributed analytic tools, analyzed data and wrote the paper.

H. L.: nalyzed data and wrote the paper.

P. A. N.: wrote the paper.

M. F.: wrote the paper.

Y. Q.: wrote the paper.

X. L.: analyzed data and wrote the paper.

P. A. B.: analyzed data and wrote the paper.

M. D.: analyzed data and wrote the paper.

G. E. K.: designed research, analyzed data and wrote the paper.

## Acknowledgment

This work was supported by the National Heart, Lung, and Blood Institute of the National Institute of Health under grant number R01HL154150. High Computing resources were provided by the Center for Computation and Visualization at Brown University.

